# The Arabidopsis AAC Proteins CIL and CIA2 Are Sub-functionalized Paralogs involved in Chloroplast Development

**DOI:** 10.1101/2021.02.02.429330

**Authors:** Mingjiu Li, Hannes Ruwe, Michael Melzer, Astrid Junker, Götz Hensel, Henning Tschiersch, Serena Schwenkert, Sindy Chamas, Christian Schmitz-Linneweber, Thomas Börner, Nils Stein

## Abstract

The Arabidopsis gene *Chloroplast Import Apparatus 2 (CIA2)* encodes a transcription factor that positively affects the activity of nuclear genes for chloroplast ribosomal proteins and chloroplast protein import machineries. *CIA2-like (CIL)* is the paralogous gene of *CIA2*. We generated a *cil* mutant by site-directed mutagenesis and compared it with *cia2* and *cia2cil* double mutant. Phenotype of the *cil* mutant did not differ from the wild type under our growth conditions, except faster growth and earlier time to flowering. Compared to *cia2,* the *cia2cil* mutant showed more impaired chloroplast functions and reduced amounts of plastid ribosomal RNAs. *In silico* analyses predict for CIA2 and CIL a C-terminal CCT domain and an N-terminal chloroplast transit peptide (cTP). Chloroplast (and potentially nuclear) localization was previously shown for HvCMF3 and HvCMF7, the homologs of CIA2 and CIL in barley. We observed nuclear localization of CIL after transient expression in Arabidopsis protoplasts. Surprisingly, transformation of *cia2* with *HvCMF3, HvCMF7* or with a truncated *CIA2* lacking the predicted cTP could partially rescue the pale-green phenotype of *cia2*. These data are discussed with respect to potentially overlapping functions between CIA2, CIL and their barley homologs and to the function of the putative cTPs of CIA2 and CIL.

**HIGHLIGHT:** The nucleus-localized CCT domain proteins CIA2 and CIL in Arabidopsis and the homologous chloroplast-localized HvCMF3 and HvCMF7 in barley retained partially overlapping functions in chloroplast development.

## INTRODUCTION

The development from proplastids to photosynthetically active chloroplasts in differentiating meristematic cells during leaf formation requires the concerted action of genes encoded by the nuclear and the chloroplast genome (plastome). In higher plants, the plastome contains around 100 genes, while the chloroplast proteome comprises more than 3000 proteins (Sugiura, 1995; Sun et al., 2009b). Consequently, the vast majority of chloroplast proteins are encoded in the nuclear genome and are subsequently imported into the plastids; in most cases by help of an N-terminal chloroplast transit peptide, cTP (Leister, 2003; Lee and Hwang, 2018; Nakai, 2018). Virtually all proteins required for the regulation of chloroplast development and the response of chloroplasts to environmental cues are nuclear-encoded and perform their function(s) in the plastids/chloroplasts, nucleus, cytoplasm or even both plastids and nucleus. Outside of plastids localized proteins might act, e.g., as transcriptional regulators or support protein import from the cytoplasm into these organelles. Only a limited number of nuclear encoded proteins with regulatory and/or non-metabolic function in chloroplast development have hitherto been characterized.

Recently, we identified a small class of nuclear-encoded proteins in seed plants (Li et al., 2019b; Li et al., 2019a) representing a subfamily of CMF (<u>C</u>CT <u>M</u>OTIF <u>F</u>AMILY) proteins (Cockram et al., 2012). The CCT domain [from the three Arabidopsis *(Arabidopsis thaliana)* proteins <u>C</u>ONSTANS, <u>C</u>ONSTANS-LIKE and <u>T</u>IMING OF CAB1] is found near the C-terminus of numerous proteins. As far as a function is known, CCT domain proteins are transcriptional (co-)regulators typically involved in modulating flowering time, light-induced signaling and circadian rhythm (Cockram et al., 2012). The CCT domain is described to support transport into the nucleus and protein-protein interactions (Kurup et al., 2000; Strayer et al., 2000; Robson et al., 2001). The members of the newly identified subfamily have only the single CCT domain in common with other CMF proteins, but share several conserved regions including a putative N-terminal cTP (Li et al., 2019a). Based on the three more intensively studied genes/proteins of this subfamily, we call this group of CCT domain containing proteins the AAC protein family [for: <u>A</u>LBOSTRIANS/HvAST/HvCMF7 (Li et al., 2019b), <u>A</u>LBOSTRIANS-LIKE/HvASL/HvCMF3 (Li et al., 2019a), and <u>C</u>HLOROPLAST IMPORT APPARATUS 2/CIA2 (Sun et al., 2001)] (Li et al., 2019a).

Like other well characterized CCT domain proteins, the Arabidopsis protein CIA2 is reported to act as a nuclear transcription factor. CIA2 stimulates the transcription of genes coding for components of the chloroplast protein import apparatus and for chloroplast ribosomal proteins. Mutation of *CIA2* leads to a pale-green phenotype and reduced chloroplast protein import (Sun et al., 2001; Sun et al., 2009a). In contrast to CIA2, the other two studied AAC proteins, the barley *(Hordeum vulgare)* HvCMF3 (ALBOSTRIANS-LIKE) and HvCMF7 (ALBOSTRIANS) are clearly localized in plastid/chloroplast and were also detected in the nucleus (Li et al., 2019b; Li et al., 2019a). Mutants of *HvCMF3* show a *xantha* phenotype and reduced amounts of chloroplast rRNAs, i.e., suffer from impaired chloroplast translation (Li et al., 2019a). Mutants of the ohnologous gene, *HvCMF7,* have albino and white-green striped leaves. They lack plastid ribosomes in white leaves and white leaf sectors, i.e., are unable to perform protein synthesis in plastids (Li et al., 2019b). Despite their different subcellular localization, the three proteins, CIA2, HvCMF3 and HvCMF7, play a critical role in chloroplast development and are needed for the correct functioning of chloroplast ribosomes. Thus, together with the presence of a predicted cTP (Emanuelsson et al., 1999), this might imply for all or for most AAC gene family members a role in normal chloroplast function and/or development.

Here we report on the Arabidopsis gene *CHLOROPLAST IMPORT APPARATUS 2-LIKE, CIL. CIA2* and *CIL* are, like *HvCMF3* and *HvCMF7,* ohnologous genes, i.e., originated as part of a whole genome duplication event early in the evolution of the *Brassicaceae* (Li et al., 2019a). We show that CIL is a nuclear localized protein. We induced a knock-out mutant of *CIL* by Cas9 endonuclease-site-directed mutagenesis and generated a double mutant, *cia2cil*. We compared *cil, cia2* and the double mutant *cia2cil* with respect to chlorophyll content, photosynthesis, chloroplast ultrastructure, and chloroplast rRNA accumulation and processing. The *cil* mutant did not express any visible phenotype different from wild type except a faster growth combined with earlier time of flowering. However, the double mutant *cia2cil* exhibited more severe defects in chloroplast development than the single mutant *cia2*. Genetic complementation of *cia2* indicated partially overlapping functions between the Arabidopsis and barley AAC genes *CIA2, HvCMF3 and HvCMF7*.

## RESULTS

### Generation of *CIL* Knock-out Mutants by Site-directed Mutagenesis Using Cas9 Endonuclease

In Arabidopsis, the closest homologs of HvCMF3 and HvCMF7 are CIA2 (AT5G57180) and CIL (AT4G25990). Sequence comparison of CIA2 and CIL revealed that both homologs share 60.6% amino acid identity as determined by alignment with Clustal Omega (Madeira et al., 2019) (Figure 1). HvCMF3/HvCMF7 in barley and CIA2 in Arabidopsis have proven to be required for chloroplast development as supported by the chlorophyll-deficient phenotype of their respective mutants (Sun et al., 2001; Sun et al., 2009a; Li et al., 2019b; Li et al., 2019a). *In silico* analyses show that, in addition to the CCT domain, all four homologs contain putative N-terminal chloroplast transit peptides, and also one or more nuclear localization signal(s) [prediction by ChloroP (Emanuelsson et al., 1999) and cNLS Mapper (Kosugi et al., 2009); Figure 1 and data not shown]. In order to check whether CIL also plays a role in chloroplast development, we utilized site-directed mutagenesis by RNA-guided Cas9 endonuclease to induce lesion(s) in the *CIL* gene. Four guide RNAs (gRNAs) were designed targeting three genomic regions of the first exon of *CIL* (Figure 2A). Two out of 10 T1 plantlets, AtCIL_P4_2 and AtCIL_P9_4, carried mutations at either or both PS2 and PS3 target sites and had chimeric genotypes (Figure 2B & Supplemental Figure 1). During propagation of the T2 progeny, a homozygous mutant, AtCIL_P4_2_18, carrying a 1 bp insertion leading to a frame shift, was selected (Figure 2C). Homozygosity of the *CIL* locus of AtCIL_P4_2_18 was confirmed by testing the T3 progeny (Figure 2D). In addition, one homogeneously biallelic mutant, AtCIL_P4_2_2_5, was identified (Figure 2D). The truncated gene in AtCIL_P4_2_18 putatively carries the information for only the N-terminal 86 in-frame amino acids of CIL, strongly suggesting that it represents a null allele, i.e., has no functional product (Figure 2E). In the following, the plants carrying the 1 bp insertion (AtCIL_P4_2_18 and corresponding T3 progenies) are referred to as *cil* mutant.

**Figure 1.**
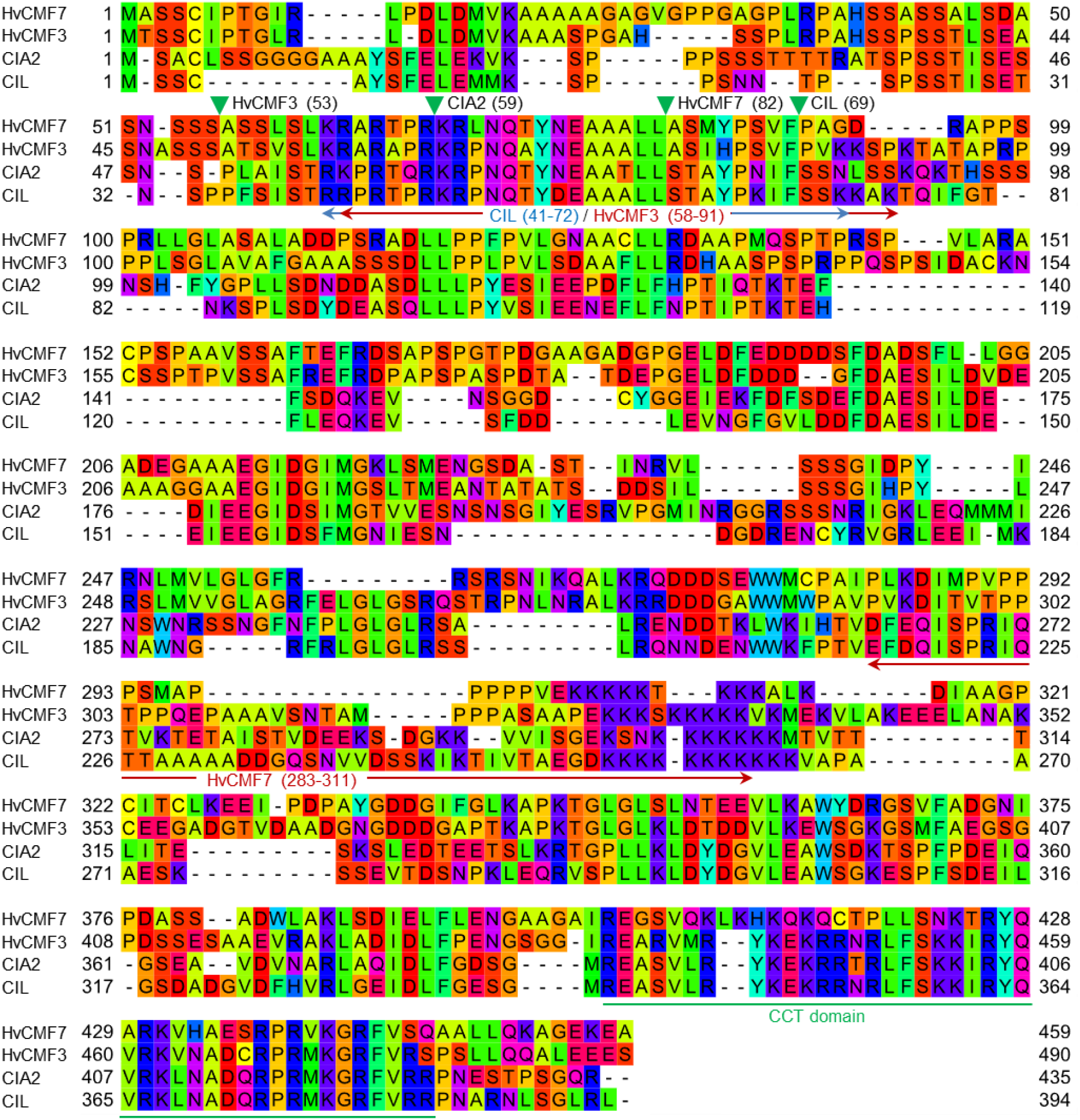
Alignment of protein sequences of HvCMF7, HvCMF3, CIA2 and CIL. Green filled triangles show cleavage sites of cTP predicted by ChloroP (Emanuelsson et al., 1999) and arrows indicate NLS (cutoff score ≥ 6.0) predicted by cNLS Mapper (Kosugi et al., 2009). Numbers in parentheses indicate amino acid positions in the respective proteins. The CCT domain is underlined in green.

**Figure 2.**
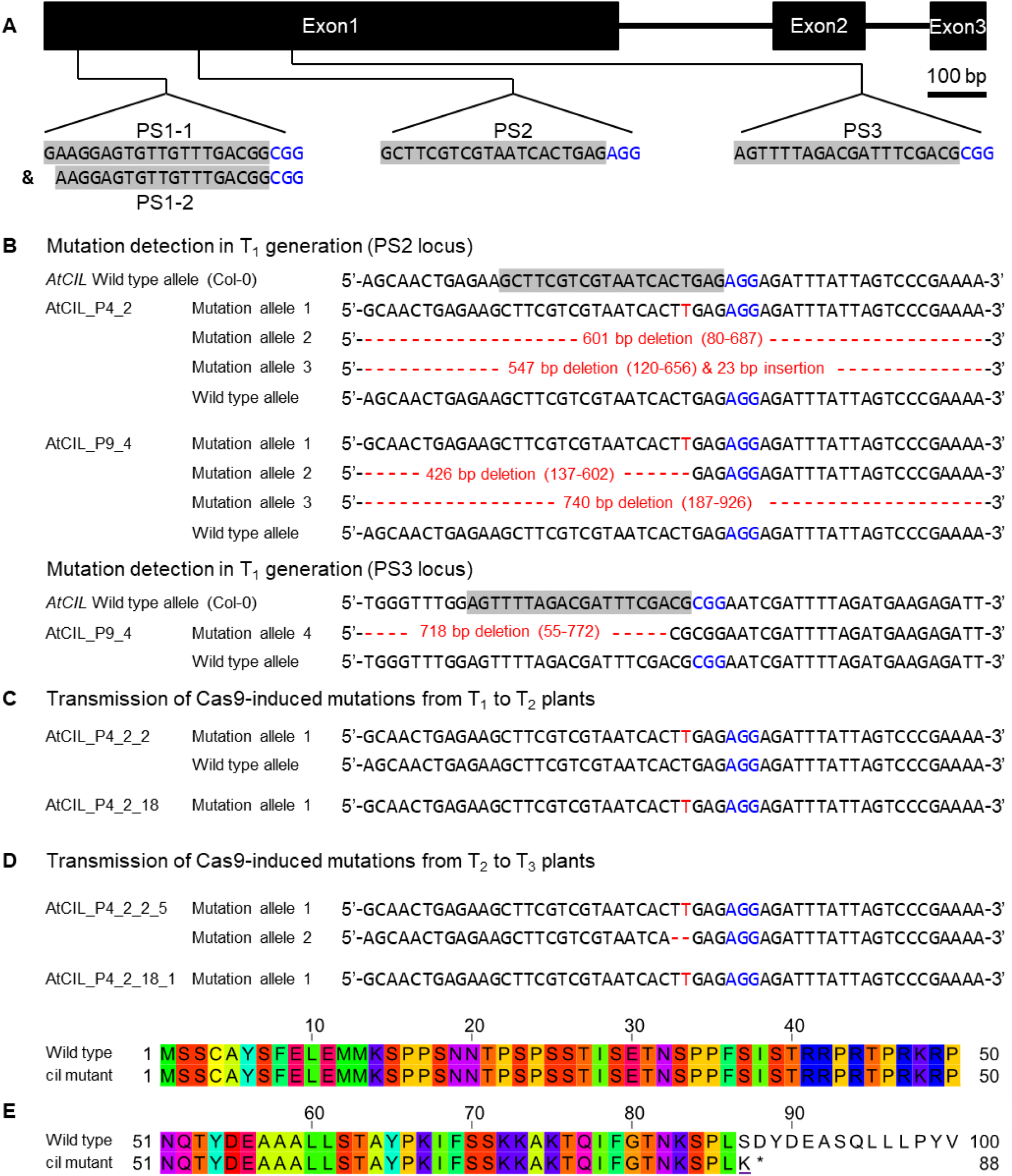
Site-directed mutagenesis of the *cil* gene by CRISPR. (A) Structure of the *cil* gene and guide RNA selection. The protospacer sequences are indicated by gray background and the protospacer adjacent motifs are highlighted by blue color. (B) Mutation detection in T_1_ plants. Two chimeric plants, AtCIL_P4_2 and AtCIL_P9_4, carrying mutations at either or both PS2 and PS3 target loci. Large deletions/insertions are represented by red dashes and small insertions by red letters. Coordinates in parentheses indicate the region of deletion. The adenine in the start codon of the *cil* sequence is counted as +1. (C) Inheritance of mutations in the T_2_ generation. The heterozygous plant AtCIL_P4_2_2 and the homozygous plant AtCIL_P4_2_18 carry a one-nucleotide insertion 3 bp upstream of the protospacer adjacent motif of the PS2 locus. (D) Inheritance of mutations in the T_3_ generation. One biallelic mutant, AtCIL_P4_2_2_5, was identified. It carries a one-nucleotide insertion and a two bp deletion. The mutation of the T_2_ plant AtCIL_P4_2_18 is stably transmitted from the T_2_ to the T_3_ generation. (E) Alignment of the deduced N-terminal protein sequences of wild type and *cil*. The altered amino acid of the mutant is underliend; an asterisk indicates the position of the immature stop codon.

### CIL Is Located in the Nucleus

*CIA2* encodes a transcription factor that activates the expression of nuclear genes, which, so far studied, code for components of the chloroplast protein translocon and for chloroplast ribosomal proteins (Sun et al., 2001; Sun et al., 2009a). Based on an increased transcription of *CIL* in *cia2*, it was proposed that CIL potentially functions as an isoform of CIA2 (Sun et al., 2001). In agreement with its proposed function as a transcriptional regulator, CIA2 is reported to be located in the nucleus (Sun et al., 2001). However, ChloroP (Emanuelsson et al., 1999) and PredSL (Petsalaki et al., 2006) predict N-terminal cTPs for CIA2 and CIL (Figure 1). As an initial step towards elucidating the molecular function of CIL, we investigated the subcellular localization of CIL by constructing a C-terminal GFP fusion to CIL, expressed under control of the Arabidopsis *Ubiquitin 10* promoter (Figure 3A). Transient expression of CIL:GFP was achieved by PEG-mediated transformation of Arabidopsis protoplasts. The green fluorescence of CIL:GFP specifically accumulated in the nucleus (Figure 3B) indicating the location of CIL in the nucleus as reported for CIA2 (Sun et al., 2001) and contrasting with the chloroplast or dual import into chloroplasts and nucleus of the barley homologs HvCMF3 and HvCMF7 (Li et al., 2019b; Li et al., 2019a).

**Figure 3.**
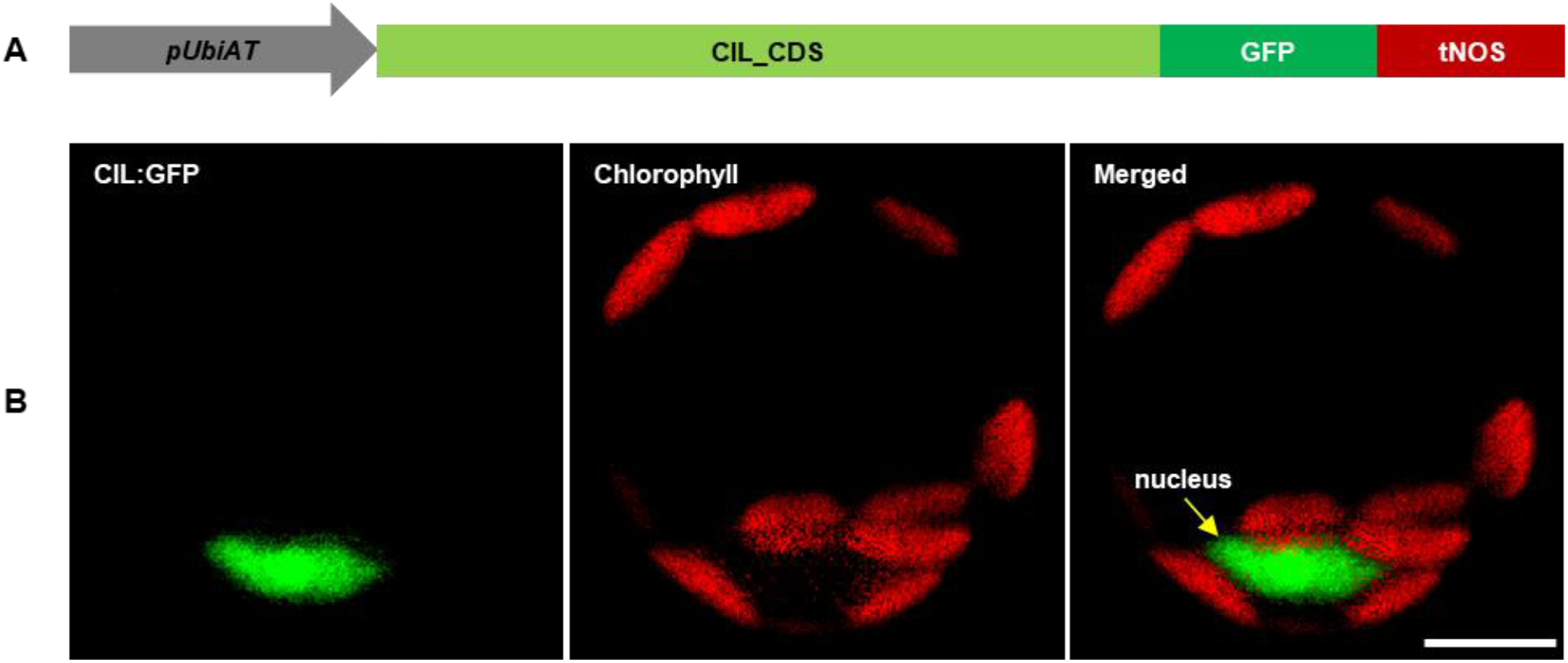
Subcellular localization of CIL:GFP fusion protein. (A) Schematic drawing of the construct used for protoplast transformation. *pUbiAT,* Arabidopsis *Ubiquitin* 10 promoter; CIL_CDS, coding sequence of the *CIL* gene; GFP, green fluorescent protein; tNOS*, Agrobacterium tumefaciens NOPALINE SYNTHASE* terminator. The drawing is not in proportion with gene length. (B) Subcellular localization of the CIL:GFP fusion protein in Arabidopsis protoplasts. Green and red colors are assigned to GFP and chlorophyll fluorescence, respectively. The yellow arrow in the merged panel indicates the nucleus. Scale bar. 10 μm.

In a further attempt to clarify whether CIL is a nuclear protein (Figure 3) or may additionally be imported into plastids, we investigated its potential import into isolated chloroplasts. Originally, the localization of CIA2 was investigated with CIA2 fused N-terminally to GUS (Sun et al., 2001), which would have masked an N-terminal cTP and made the detection of a chloroplast import of CIA2 unlikely. Therefore, we included CIA2 in the assay. CIA2 and CIL were translated and radiolabeled with [^35^S]-methionine in reticulocyte lysates and incubated with isolated pea (*Pisum sativum*) chloroplasts. As a positive control the stromal protein FERREDOXIN-NADP(+) REDUCTASE (FNR) was used (Guan et al., 2019). FNR was imported as expected, since we observed the mature, processed form of the protein after incubation with chloroplasts. The mature protein was resistant to thermolysin treatment showing that FNR was inside of intact and import competent chloroplasts (Supplemental Figure 2). In contrast, only faint bands were visible after the import reaction and removal of the translation product in case of CIA2 and CIL. Moreover, none of these bands was resistant to thermolysin treatment indicating that both proteins were not transported into chloroplasts (Supplemental Figure 2). Therefore, CIA2 and CIL represent members of the AAC subfamily that are located in the nucleus.

### Chlorophyll Content and Photosynthetic Parameters of *cia2, cil* and Double Mutant *cia2cil*

The *cia2* mutant exhibits a pale-green phenotype (Figure 4A) as previously described by Sun et al. (2001), while the *cil* mutant did not differ phenotypically from the wild type (Col-0) under the growth conditions used in this study (Figure 4A, D-F). We generated a *cia2cil* double mutant by crossing the original EMS-induced *cia2* mutant with the newly generated *cil* mutant (AtCIL_P4_2_18_1). All obtained 13 F1 hybrids showed a normal green phenotype. The homozygous *cia2cil* double mutants of the F2 generation, however, exhibited a distinctly retarded growth compared to *cia2*, *cil* and the wild type (Supplemental Figure 3F) and a more severe chlorophyll-deficient phenotype than *cia2* (Figure 4A-F; Supplemental Figure 3G) confirmed by measurements of the chlorophyll *a* and *b* contents (Figure 4D-E). The double mutant started flowering three days later than wild type and *cia2,* while *cil,* interestingly, started flowering two days earlier than wild type and *cia2* (Supplemental Table 1). The earlier flowering of the *cil* mutant correlates with its faster growth as reflected by the projected leaf area (Supplemental Figure 3F). Similar to wild type Col-0, all single and double mutants developed 14 rosette leaves until reaching the bolting stage. Therefore, the differences between these lines with respect to day to flowering are may be explained by physiological changes that leading to different growth rates. The double mutant shows a delayed greening, i.e., the chlorophyll deficiency of *cia2cil* is more pronounced in young leaves. The green leaf pigmentation is increasing with further development; however, the leaves remain paler than in wild type (Figure 4A; Supplemental Figures 3G & 4A). The *cia2* single and *cia2cil* double mutants exhibited a higher chlorophyll *a:b* ratio than wild type (Figure 4F). Since photosystem II (PSII) is enriched in chlorophyll *b* as compared to PSI, the higher chlorophyll *a:b* ratio could be an indication that PSII is more severely affected than PSI in *cia2* and *cia2cil* mutants.

**Figure 4.**
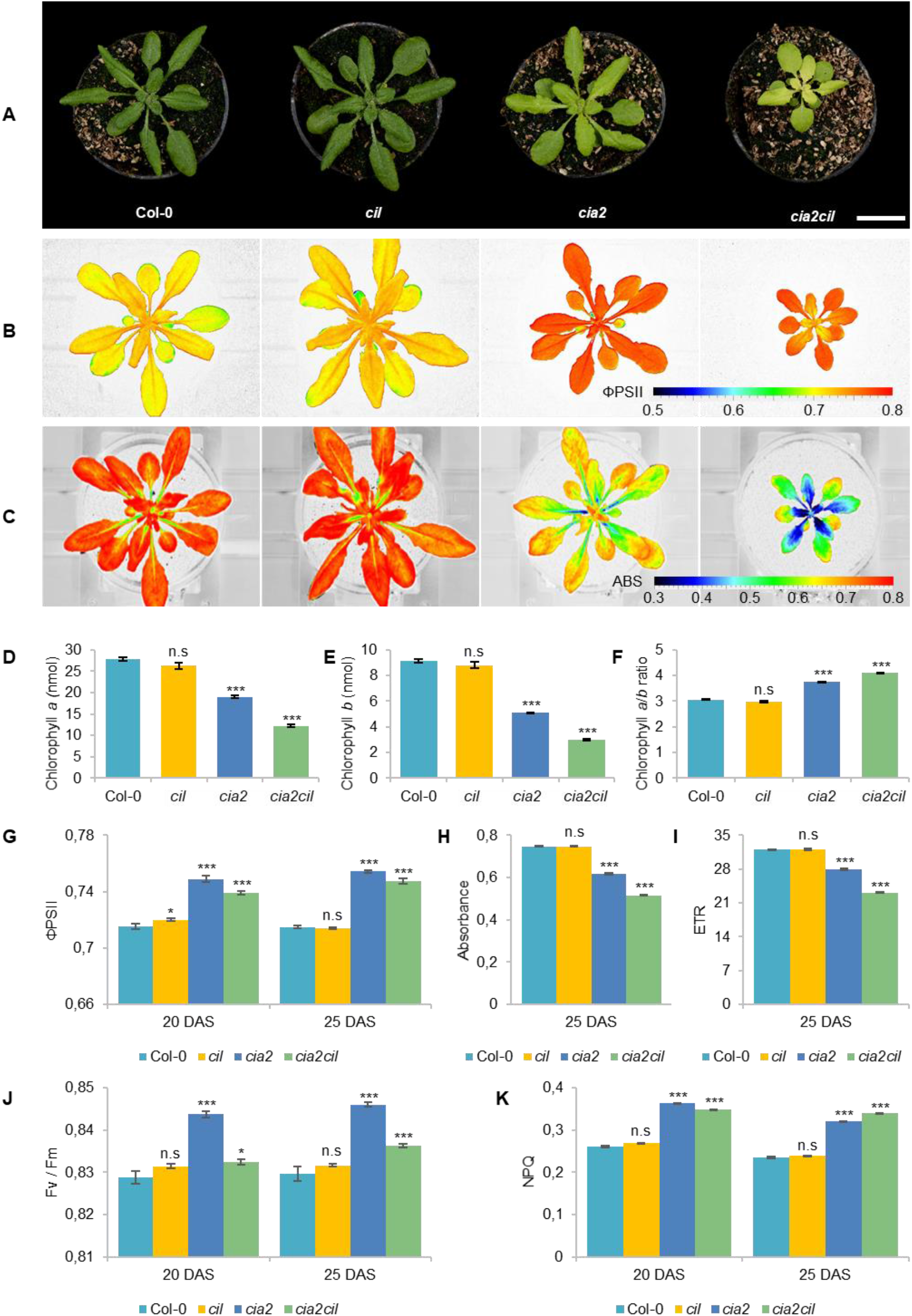
Measurement of chlorophyll content and photosynthetic performance of wild type Col-0 and *cil, cia2* and *cia2cil* mutants. (A) Phenotype of Col-0, *cil, cia2* and *cia2cil* plants. Scale bar, 2 cm. (B) False-color images of the operating light use efficiency of PSII (ΦPSII). (C) False-color images of NDVI imaging of actinic light absorbance. (D-F) Quantification of chlorophyll *a* (D), chlorophyll *b* (E) and ratios of chlorophyll *a* to *b* (F). (G-K) Measurement of photosynthetic parameters. Results are presented as mean ± SEM. Number of plants used for chlorophyll content measurement N = 10; number of plants used for photosynthetic measurement N = 15. *Student’s t-test* (Tails = 2; Type = 2) significant levels, n.s, not significant, * *p* < 0.05, * * *p* < 0.01, * * * *p* < 0.001. ΦPSII, photosystem II operating efficiency; Absorbance, absorbance of actinic light; ETR, electron transport rate; F_v_/F_m_, maximum quantum yield of PSII photochemistry measured in the dark-adapted state; NPQ, non-photochemical quenching. DAS, days after sowing. Plants and images shown in panels A-C were at developmental stages 20 DAS. All photosynthetic measurements were performed at 120 μE actinic light.

We used non-invasive chlorophyll fluorescence imaging integrated into an automated, conveyor-based phenotyping platform to quantify photosynthesis-related traits of *cil, cia2* and *cia2cil* (Junker et al., 2015; Tschiersch et al., 2017). The PSII operating efficiency (ΦPSII) of *cil* did not differ from Col-0. In contrast, the respective levels were mildly but significantly increased in *cia2* and *cia2cil* compared to Col-0 (Figure 4B & 4G). The elevation of ΦPSII in *cia2* and *cia2cil* was independent of light intensity and light-/dark-adaptation; as expected, ΦPSII values decreased at high light intensity of 400 μE as compared to low light intensity of 120 μE (Figure 4B & 4G; Supplemental Figure 3A-C). Intriguingly, as revealed by NDVI (normalized difference vegetation index) imaging, *cia2* and *cia2cil*, compared to the wild type and *cil*, showed a substantial decrease in the absorbance of actinic light (Figure 4C & 4H) resulting in a lower electron transport rate (Figure 4I and Supplemental Figure 3E). This suggests that the slightly increased PSII efficiency of *cia2* and *cia2cil* cannot compensate for the low amount of energy absorbed by PSII under steady-state light conditions. Next, we examined the role of photochemical quenching in *cil, cia2* and *cia2cil* by measuring the maximum quantum yield of PSII photochemistry in the dark-adapted state (F_v_/F_m_), the ‘excess excitation energy’ indicator non-photochemical quenching (NPQ) and PSII efficiency factor qP (representing the fraction of open PSII reaction centers). There was no difference between *cil* and Col-0 for all measured parameters. In line with the observed higher PSII operating efficiency, the F_v_/F_m_ values were significantly higher in *cia2* and *cia2cil*. Interestingly, *cia2* exhibited a higher F_v_/F_m_ than the *cia2cil* double mutant (Figure 4J). Also, the NPQ and qP values were significantly increased in both mutants (Figure 4K; Supplemental Figure 3D). The increase of F_v_/F_m_ in the dark-adapted state could indicate that *cia2* and *cia2cil* have intrinsically more efficient PSII reaction centers, whereas the increase in photochemical quenching agrees with a larger fraction of PSII reaction centers performing photochemistry in *cia2* and *cia2cil*.

Overall, these results demonstrate that the mutations in *cia2* and *cia2cil* do not affect the primary function of PSII as indicated by the high values of maximum quantum yield of PSII (F_v_/F_m_). The decreased photosynthetic activity of the *cia2* and *cia2cil* mutants is due to the substantially decreased absorbance of actinic light, resulting in a lower electron transport rate. Moreover, mutation of the *CIL* gene alone has no significant effect on photosynthetic performance.

### CIA2 and CIL Affect Thylakoid Organization

The observed chlorophyll deficiencies and differences in photosynthetic parameters between *cia2, cia2cil* double mutant and wild type suggest structural defects of chloroplasts caused by mutation. Therefore, we examined and compared chloroplast ultrastructure in wild type and mutants by transmission electron microscopy. Mutation of *CIA2* and/or *CIL* alters chloroplast morphology and the internal organization of the photosynthetic apparatus. Compared to Col-0, chloroplast size (width and length) was reduced in *cia2* and *cia2cil* mutants while *cil* mutants had smaller chloroplasts with reduced width only (Figure 5A-K). Further, the structure of grana was quantified within 1 μm^2^ areas. We observed a significant increase in the number of grana in *cil, cia2* and *cia2cil* compared to wild type (Figure 5L). A higher number of thylakoid membranes within each granum was observed in *cil,* but this number was lower in *cia2cil* and remained at the same level in *cia2* as compared to Col-0 (Figure 5M). The distance between each thylakoid was not different between wild type and mutants (Figure 5N). Finally, we measured the maximal height of grana in 40-80 chloroplasts of wild type and mutants. Compared to Col-0, the *cil* mutant contained grana composed of a higher number of thylakoid membranes; on the contrary, the maximal height of the grana and the number of thylakoids in the largest grana were significantly reduced in both *cia2* and *cia2cil* (Figure 5O-P). The lower number of thylakoids per granum is most likely responsible for the observed lower light absorbance and lower electron transport rate of the *cia2* and *cia2cil* mutants.

**Figure 5.**
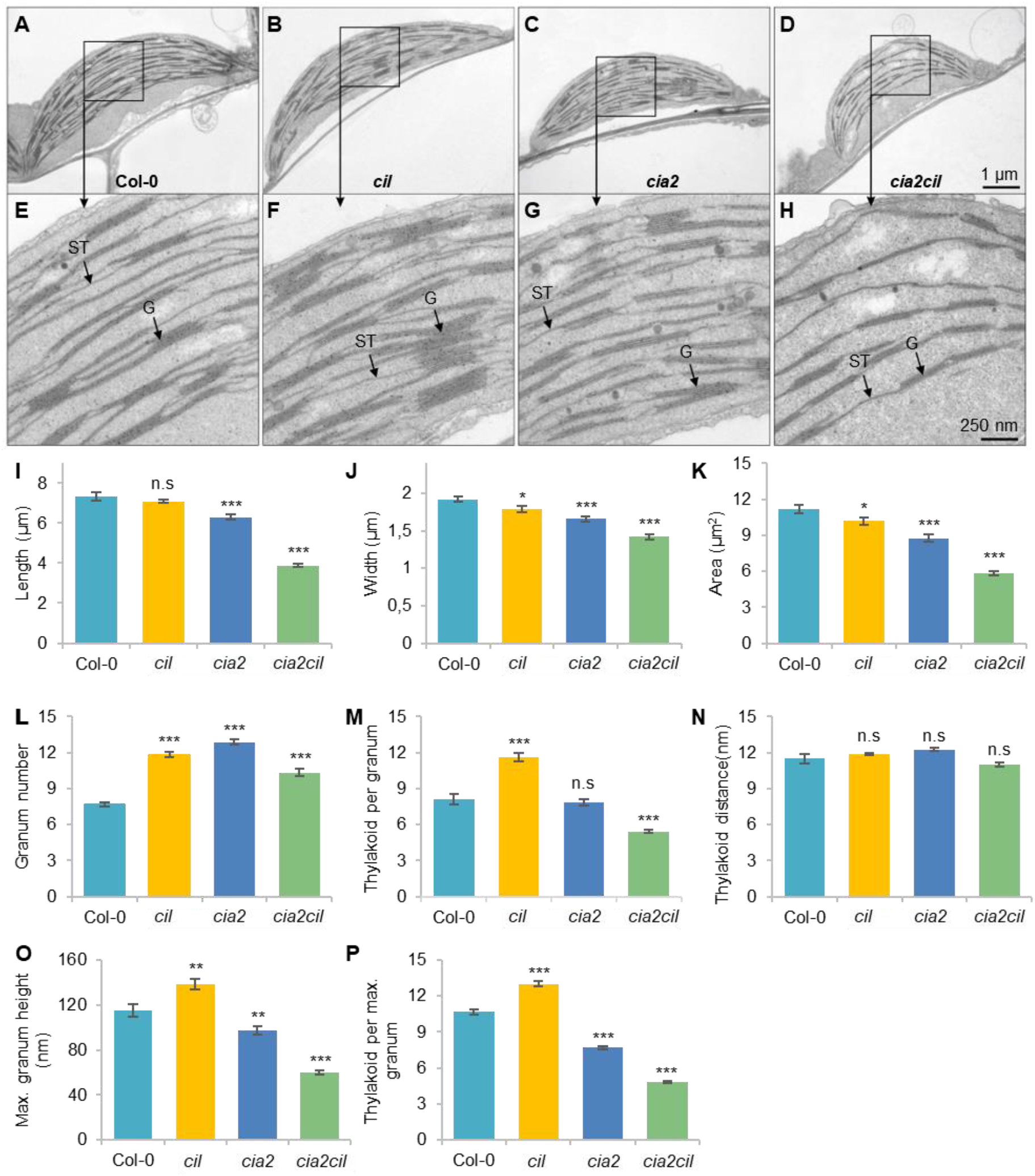
Ultrastructural analysis of chloroplasts of the wild type Col-0, and *cia2, cil,* and *cia2cil* mutants. (A-H) Comparison of the chloroplast ultrastructure in wild type and mutants. Panels E to H represent larger magnification of marked areas of the corresponding plastid in the top panels A to D, respectively. ST, stroma thylakoid; G, granum. (I-P) Quantification of chloroplast architecture and components of photosynthetic apparatus. The granum number (L) and thylakoid per granum (M) were counted within an area of 1 μm^2^. Max. granum height (O) represents the granum with maximum number of thylakoid membranes within each chloroplast. Thylakoid per max. granum (P) represents the number of thylakoid membranes within the highest granum. Results are presented as mean ± SEM (N ≥ 40). *Student’s t-test* (Tails = 2; Type = 3) significant levels, n.s, not significant, * *p* < 0.05, * * *p* < 0.01, * * * *p* < 0.001.

### Impaired Chloroplast rRNA Processing in *cia2* and *cia2cil*

Microarray expression analysis revealed a down-regulation of genes for chloroplast ribosomal proteins in *cia2* suggesting a role of CIA2 in maintaining chloroplast translation efficiency (Sun et al., 2009a). Since rRNAs do not accumulate if not incorporated into ribosomal subunits, the abundance of individual rRNA species serves as a proxy for the accumulation of the respective ribosomal subunits (Fristedt et al., 2014). Separation of cytosolic and chloroplast rRNA species on agarose gels revealed no striking difference between Col-0 and the *cil, cia2* as well as *cia2cil* mutants in mature leaves (Figure 6B; Supplemental Figure 4). While cytosolic and chloroplast rRNAs did apparently not differ in their abundance in young leaves among Col-0, *cil* and *cia2,* the amounts of the chloroplast rRNAs were distinctly reduced in young leaves of *cia2cil,* the 16S rRNA to lesser extent than the 23S rRNA (Figure 6B). Notably, the 2.9 kb and 2.4 kb 23S rRNA precursors were faintly visible in *cia2* and *cia2cil,* suggesting defects in rRNA processing in both mutants (Figure 6A-B). RNA gel-blot analysis with specific probes against the 16S and 23S rRNAs revealed an inefficient processing of the 2.9 kb and 2.4 kb 23S rRNA precursors in both mature and young leaves of *cia2* and *cia2cil,* with the young leaves being more severely affected (Figure 6C). We observed also an over-accumulation of the unprocessed 1.7 kb 16S rRNA precursor in young leaves of the *cia2cil* mutant, suggesting inefficient 3’ trimming of the 1.9 kb 16S precursor (Figure 6A & 6D; Supplemental Figure 5). Thus, we obtained evidence for impaired chloroplast rRNA processing in *cia2* and, to much more extent, in *cia2cil,* whereas *cil* did not differ from wild type.

**Figure 6.**
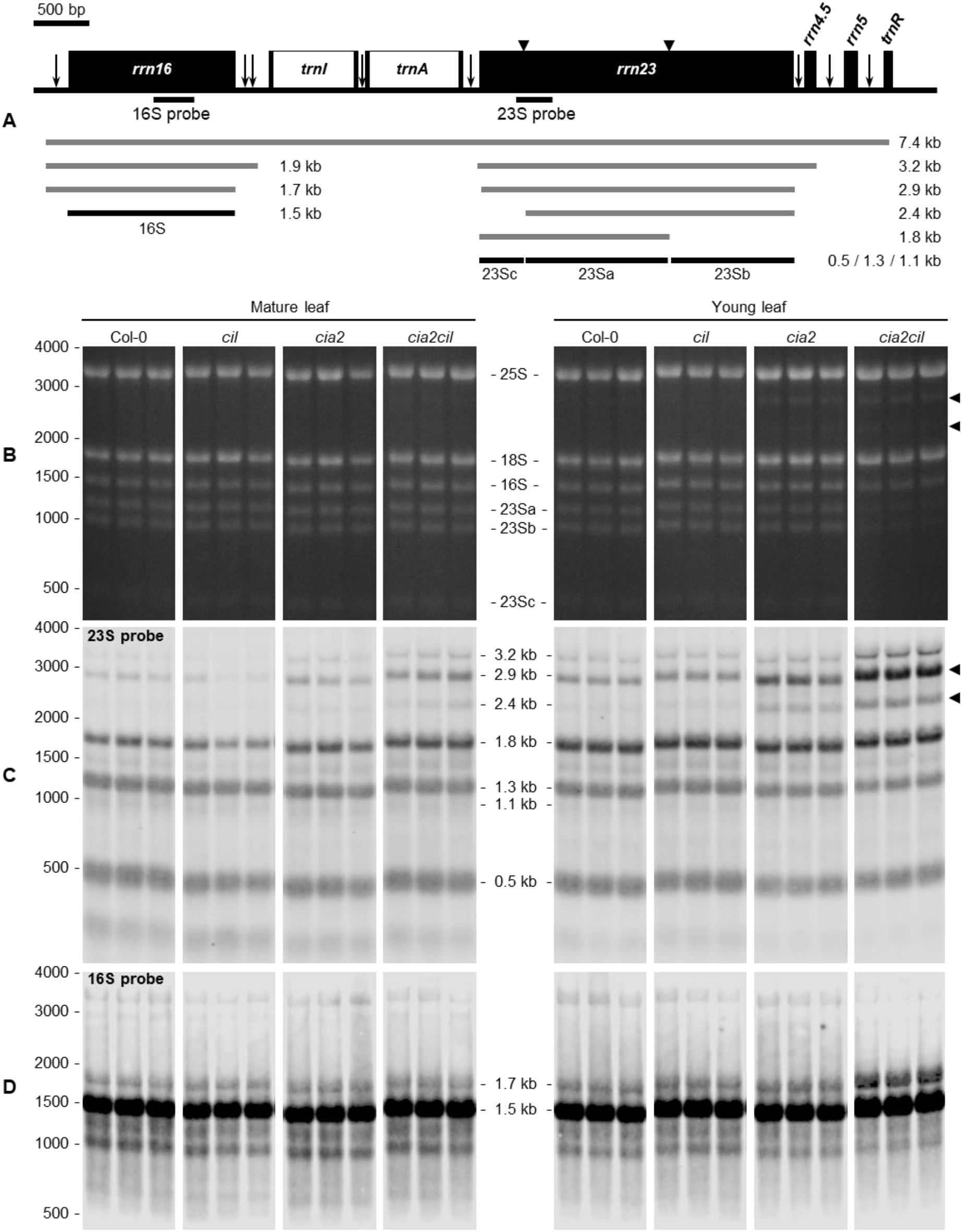
Analysis of chloroplast rRNA processing in Col-0, *cil, cia2* and *cia2cil* mutant plants. (A) Structure and transcript pattern of the chloroplast *rrn* operon. Black boxes indicate exons and white boxes indicate introns. Vertical arrows indicate processing sites in the primary transcripts of the *rrn* operon. Positions of internal cleavage sites (hidden breaks) in the 23S rRNA are shown as black triangles. Positions of the hybridization probes for 16S and 23S are indicated below the operon structure. The 7.4-kb primary transcript and various processing precursors are shown with grey lines; the mature forms of 16S and 23S rRNAs are shown with black lines. The Arabidopsis chloroplast genome (GenBank accession number: NC000932.1) was used as reference. (B) Separation of cytosolic and chloroplastic rRNAs on agarose gel. The 23Sa, 23Sb and 23Sc bands represent 1.3 kb, 1.1 kb and 0.5 kb mature forms of 23S rRNA, respectively. Each sample with 3 biological replicates. Arrows indicate intermediates of inefficient processing of the 23S rRNA. (C) Analysis of 23S rRNA processing by RNA gel-blot hybridization. Arrows indicate pre-mature 23S rRNA species shown in panel B. (D) Analysis of 16S rRNA processing by RNA gel-blot hybridization. The original gel images are provided as Supplemental Figure 4.

### Barley Homologs and N-Terminally Truncated CIA2 Improve Chlorophyll Content in *cia2*

*Hvcmf3, Hvcmf7, cia2* and *cil* (in the double mutant *cia2cil)* caused chloroplast ribosome deficiencies despite different subcellular localization of their gene products. A plausible assumption is that the barley and Arabidopsis genes might have overlapping functions. As *cil* lacks a visible phenotype and in order to rule out the additive effect of *cil* in the *cia2cil* mutant, we attempted heterologous complementation of *cia2* by *HvCMF3* and *HvCMF7,* respectively. Both barley genes were able to improve the *cia2* pale-green phenotype; the chlorophyll contents of the complementation lines were mildly but significantly increased compared to *cia2* (Figure 7A-B & 7D).

**Figure 7.**
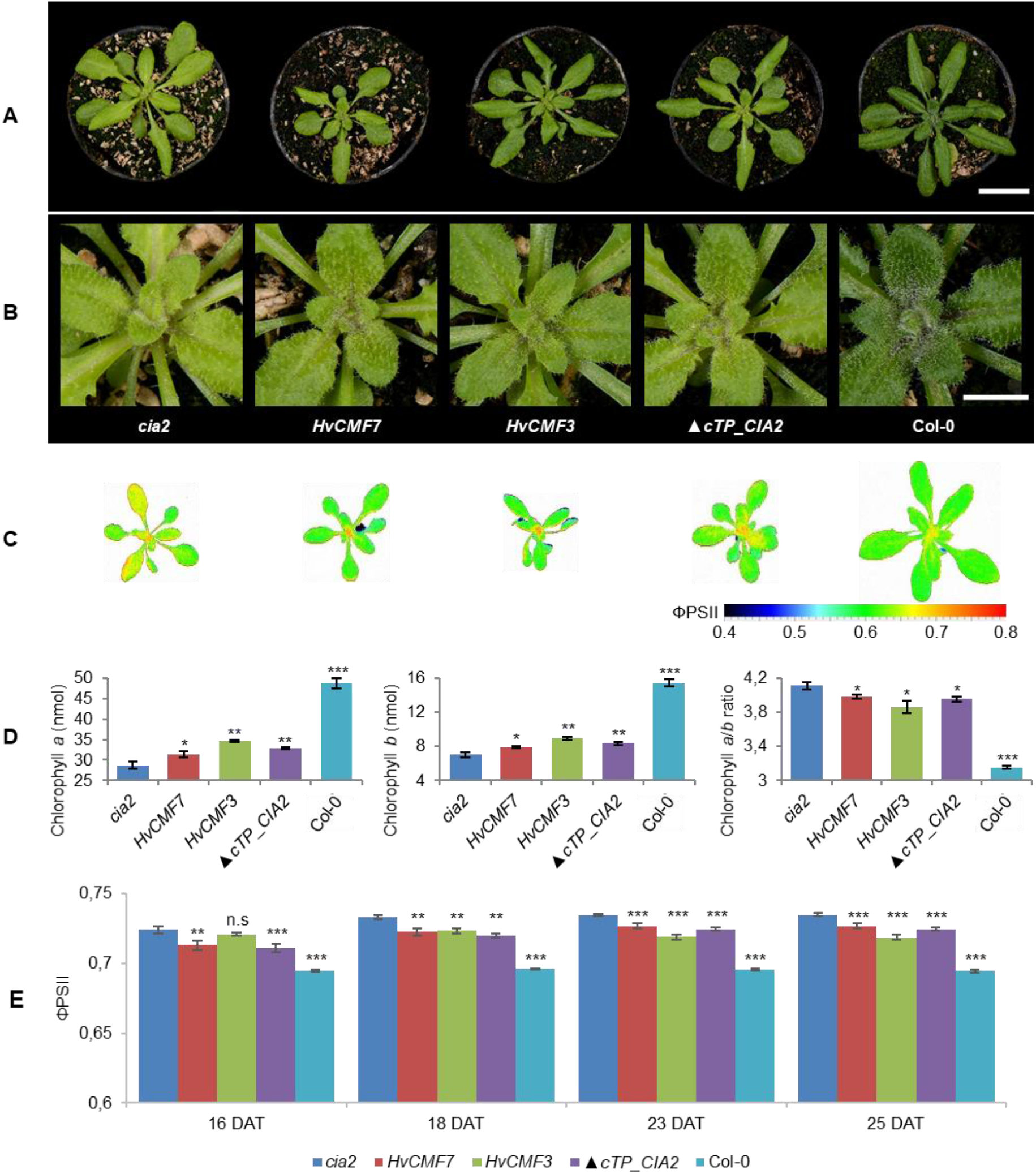
Genetic complementation of the *cia2* mutant with *HvCMF3, HvCMF7* and ▲ *cTP_CIA2*. (A) Phenotype of *cia2* mutant, complementation lines and wild type. (B) Enlargement of the central region of the respective plants in panel A showing phenotype of the young vegetative rosette leaves. (C) False-color images of the operating light use efficiency of PSII (ΦPSII). (D) Quantification of chlorophyll contents. (E) Quantification of photosynthesis operating efficiency. Results are presented as mean ± SEM (N ≥ 15 in panel D; N ≥ 9 in panel E). *Student’s t-test* (Tails =2; Type = 3) significant levels, n.s, not significant, * *p* < 0.05, * * *p* < 0.01, * * * *p* < 0.001. DAT, days after transfer to soil. Scale bar in panel A, 2 cm. Scale bar in panel B, 0.5 cm.

ChloroP (Emanuelsson et al., 1999) predicts for the N-terminal 59 amino acids of CIA2 to function as chloroplast transit peptide (cTP). CIA2 was localized to the nucleus and not to the plastids in a previous study (Sun et al., 2001). However, GUS was fused N-terminally to CIA2 in this study thus hiding the putative cTP, i.e., this experiment did not exclude a potential chloroplast localization of CIA2. Yet, CIA2 was targeted only to the nucleus also in the present investigation. To test whether the predicted cTP is essential for CIA2 protein activity, we attempted to complement the *cia2* mutant with a truncated form of CIA2 (▲ *cTP_CIA2;* CIA2 without the predicted N-terminal cTP). Very similar to the phenotype of the *HvCMF3/HvCMF7* heterologous complementation lines, CIA2 without cTP was able to rescue partially the pale-green phenotype (Figure 7A-B & 7D). The chlorophyll content of transgenic plants bearing the empty vector control without insertion of (▲ *cTP_CIA2, CIL, HvCMF3* or *HvCMF7*) remained at the level observed with the *cia2* mutant. Therefore, the enhanced chlorophyll content of the complementation lines most likely results from the expression of the transgene.

To further characterize the transgenic lines, the PSII operating efficiency (ΦPSII) of light-adapted plants was measured at 16, 18, 23 and 25 DAT (DAT, days after transfer to soil). Compared to *cia2*, the transgenic lines showed significantly lower ΦPSII values, i.e., got closer to the level of Col-0 (Figure 7C & 7E). As in case of chlorophyll content, the *HvCMF3* transgenic line showed the largest effect compared to plants derived from the other two transgenic lines, observed at time points 23 and 25 DAT (Figure 7E). Quenching analysis of dark-adapted plants indicated a slight increase of the F_v_/F_m_ value of the *HvCMF7* complementation line at 23 and 25 DAT, but not of the *HvCMF3* complementation lines. ▲ *cTP_CIA2* complementation lines exhibited a slight increase at 23 DAT, but dropped back to the *cia2* level at 25 DAT (Supplemental Figure 6A). Non-photochemical quenching (NPQ) and photochemical quenching (qP) were increased in *cia2* compared to wild type (Figure 4K and Supplemental Figure 3D). Complementation of *cia2* with *HvCMF3* or *HvCMF7* but not with ▲ *cTP_CIA2* (except at time point 23 DAT for the qP value) reduced the mean values of NPQ and qP. Again, complementation by *HvCMF3* led to the largest degree of decrease (Supplemental Figure 6B-C). Taken together, the analyses of chlorophyll content and of photosynthetic parameters indicate a partial reduction of the effects of the *CIA2* mutation by complementation with *HvCMF3, HvCMF7* and ▲ *cTP_CIA2*.

## DISCUSSION

Transition of plastids into photosynthetic active chloroplasts requires the concerted action of the plastome and the nuclear genome. According to first studies on AtCIA2, HVCMF3 (ALBOSTRIANS-LIKE) and HvCMF7 (ALBOSTRIANS), the members of the small AAC subfamily of CCT motif proteins belong to nucleus encoded proteins that play essential roles in chloroplast development (Sun et al., 2001; Li et al., 2019b; Li et al., 2019a). Our present results indicate that the Arabidopsis AAC protein CIL has very similar functions to its ohnolog CIA2. Moreover, our data suggests partial functional coincidence of the barley proteins CMF3 and CMF7 with CIA2 although the barley and Arabidopsis proteins differ with respect to their subcellular localization.

As a first step towards elucidating the function of CIL, we generated a putative null mutant of the *CIL* gene. Because of the pale-green phenotype of *cia2* and the reported function of CIA2 as a transcriptional regulator of genes for chloroplast protein transport and chloroplast ribosomal proteins (Sun et al., 2001; Sun et al., 2009a), the majority of our investigation concerned parameters of *cil,* which are related to chloroplast function and development. To our surprise, we observed no significant differences between *cil* and wild type with respect to chlorophyll content, photosynthetic parameters, amount of chloroplast rRNA and chloroplast rRNA processing. These results and similar observations recently published by Gawronski et al. (2020) suggest that CIA2 and CIL have at least partially redundant functions and CIA2 activity alone is sufficient for growth and development under normal growth conditions. This conclusion is supported by the distinctly higher expression level (log2 fold-change > 2) of *CIA2* compared to *CIL* in Col-0 according to the eFP expression atlas. Otherwise, both genes have a similar expression pattern. They are expressed in all organs and all developmental stages except mature siliques and roots of mature plants (Supplemental Figure 7; Arabidopsis eFP Browser, http://bar.utoronto.ca/) (Sun et al., 2001; Winter et al., 2007; Hruz et al., 2008).

The comparison of *cil* and *cia2* with their double mutant *cia2cil* further corroborates the hypothesis of redundant functions of CIL and CIA2. Although the visible phenotype and most other analyzed parameters of *cil* did not differ from Col-0, the double mutant showed more severe differences to the wild type than *cia2*. Obviously, mutation of *cil* potentiates the effects of the *cia2* mutation. This phenomenon is called ‘phenotype gap’ (Ewen-Campen et al., 2017). It refers to the fact that, due to functional redundancy, knock out of one gene in a pair of paralogs leads to a loss-of-function mutant without a detectable phenotype. In Arabidopsis, a compilation of 70 paralogous gene pairs with loss-of-function in one paralog resembles the case of *CIA2/CIL* (Lloyd and Meinke, 2012), i.e., the mutant of only one of the two paralogous genes showed a phenotype, which was more severely expressed in the double mutant. The fact, that *CIA2* and *CIL* have been kept as active genes in the genome since their occurrence as result of the whole genome duplication in the *Brassicaceae* (Muhlhausen and Kollmar, 2013), makes sub-functionalization (Lynch and Conery, 2000) of two homologous genes likely. The pattern of gene expression over most stages of development in green as well as non-green tissues suggests that CIL and CIA2 are not exclusively involved in chloroplast biogenesis. A first hint supporting this assumption may be our observation of faster growth and earlier flowering of *cil* compared to Col-0 and the other analyzed mutants. Yang and Sun (2020) reported on interactions of CIA2 and CIL with flowering-control proteins. Our data points to a more general effect of CIL on development.

The impaired processing of plastid 16S and 23S rRNAs observed in *cia2cil* and to lesser extent in *cia2* may explain the reduced amount of plastid rRNAs in *cia2cil* and may negatively affect ribosome function, i.e., protein synthesis in chloroplasts. Chloroplast rRNA processing requires several specific enzymes and the assembly of the precursor rRNA with ribosomal proteins (e.g., Barkan, 1993; Yu et al., 2008; Jiang et al., 2018). Thus, the function proposed for CIA2 by Sun et al. (2009a) as transcription regulator of genes coding for chloroplast ribosomal proteins and for components of the translocon required for protein import into chloroplasts, is in agreement with the observed delayed chloroplast rRNA processing in *cia2* and *cia2cil* and reduced amounts of plastid rRNAs in *cia2cil*. The more serious effect of the two mutations in *cia2cil* compared to *cia2* on 16S and 23SrRNA processing, chlorophyll content, thylakoid/grana structure and, in particular, photosynthetic parameters may be responsible for the slower growth of *cia2cil vs. cia2, cil* and wild type. Mutant analyses demonstrated that inefficient chloroplast translation is often associated with altered organization of the thylakoid membrane system (Fristedt et al., 2014; Liu et al., 2015; Zhang et al., 2017; Li et al., 2019a). The delayed greening phenotype of *cia2cil* resembles the *rbf1-1* mutant with defects in chloroplast translation due to inefficient processing of 16S rRNA (Fristedt et al., 2014). Chloroplast development has an extraordinarily high demand for *de novo* protein biosynthesis. Therefore, even a mild disturbance of the translational machinery of the plastids may result in delayed greening of the young developing leaves. The demand for translation capacity decreases during the subsequent aging process and the translational machinery in mutants with a mild plastid ribosome deficiency gets time to catch up and restores to wild type levels of chloroplast proteins (Fristedt et al., 2014). Impaired import of proteins into chloroplasts as reported for CIA2 (Sun et al., 2001) could also directly affect the structure of thylakoids, the function of photosynthesis and the synthesis of chlorophylls since virtually all functions of chloroplasts need nuclear-gene-encoded proteins imported from the cytoplasm (Jarvis et al., 1998; Bauer et al., 2000).

A nuclear localization of CIL would be a precondition to function like its homolog CIA2 as a transcription regulator of nuclear genes involved in chloroplast development. A suite of programs is available to predict *in silico* the subcellular localization of proteins. Interestingly, nuclear localization signals, NLS, but also chloroplast transit peptides, cTPs, are predicted for CIA2 and CIL as well as for their barley homologs HvCMF3 and HvCMF7. In correspondence with the predictions, we detected HvCMF3 and HVCMF7 in chloroplasts and, according to preliminary data, in the nucleus (Li et al., 2019b; Li et al., 2019a). In our present study, we detected CIL, C-terminally fused with GFP, only in the nucleus. There was no indication for a transport into chloroplasts or other compartments of Arabidopsis protoplasts. Moreover, neither CIL nor CIA2 were imported into isolated chloroplasts. In contrast to the situation with FNR, neither the imported processed proteins nor the preproteins of CIA2 and CIL were observed after the import assays (Supplemental Figure 2, lanes 2 and 3). Thus, the *in silico* predicted cTPs (Figure 1) [e.g., by ChloroP (Emanuelsson et al., 1999) and PredSL (Petsalaki et al., 2006)] do not seem to support the binding to the envelop of the chloroplasts and not the import of the two Arabidopsis proteins into these organelles (Supplemental Figure 2). Gawronski et al. (2020) recently reported also an import of CIL exclusively into the nucleus; however, they observed an import of CIA2 into both plastids and nucleus. They used experimental conditions that differed from those in our experiments, e.g., expression of CIA2 by a different promoter. More studies are needed to find out whether the import of CIA2 takes place only under certain conditions. We have, for example, not checked the possibility of a role in the transport of proteins into plastid types other than the investigated chloroplasts, e.g., into plastids at an earlier stage in their development to chloroplasts. The conservation of the N-terminal amino acid sequence extending over the predicted cTP (Figure 1) suggests, however, that this part of the AAC proteins is of functional importance. *In silico* analyses suggest a different (or additional to cTP) function of the N-terminal region. The tools WoLF PSORT (Horton et al., 2007) and Localizer (Sperschneider et al., 2017) predict a high probability for a nuclear localization of CIA2 and CIL since they possess potential NLS near their N-terminus and in other regions. NLS function is also predicted for the N-terminal regions of HvCMF3 and HvCMF7 and supported by the observation of nuclear localization of GFP, N-terminally fused with the predicted cTPs of the two barley proteins (Li et al., 2019b; Li et al., 2019a). Yang and Sun (2020) reported functional NLS in the N-terminal regions of CIA2 and CIL.

A function of the N-terminal domain additional to or other than as cTP is also suggested by results of our attempts to rescue the *cia2* mutant by transformation with a construct expressing a truncated form of CIA2 that lacks completely the predicted N-terminal cTP and NLS (Figure 1). The truncated CIA2 should not be imported into chloroplasts. However, it might still enter the nucleus by diffusion or with support of the predicted NLS other than the NLS near the N-terminus. The size of CIL makes the necessity of an active transport into the nucleus very likely, but at least minor amounts may still diffuse into the nucleus (Wang and Brattain, 2007) and explain the observed limited rescuing of the *cia2* mutation. Alternatively, the rather low rescue potency of the truncated protein could be an indication for the importance of the N-terminal region for normal function of CIA2 in the nucleus. We conclude that CIL is a nuclear protein with low probability for a transport into chloroplasts, CIA2, CMF7 and CMF3 are most likely dually transported into plastids and nuclei, and the N-terminal region of the two Arabidopsis proteins CIL and CIA2 and of the barley proteins CMF3 and CMF7 contain NLS that support their active transport into the nucleus.

*Hvcmf3* showing a mild chloroplast ribosome deficiency and *Hvcmf7* lacking plastid ribosomes in albino tissue share defects in chloroplast translation with the Arabidopsis mutants *cia2* and *cia2cil*. Moreover, the four genes have similar structures and, although there is evidence for a chloroplast localization of HvCMF3 and HvCMF7, the two barley proteins are likely transported into both chloroplasts and nucleus (Li et al., 2019b; Li et al., 2019a). Depending on the prediction program used, sequence analyses suggest for all four proteins potential nuclear and/or chloroplast localization. Therefore, we attempted to rescue the *cia2* mutant by transformation with *HvCMF3* and *HvCMF7*. The obtained transgenic lines showed a partial rescue of the chlorophyll deficiency and improved photosynthetic performance suggesting that the barley proteins or their downstream effects may act in the same compartment and have partially similar function(s) as CIA2. Due to their conserved domain structure, e.g., the CCT domain, they might be able to substitute CIA2 in protein-protein interactions, though with lower efficiency.

In our present study we characterized for the first time a mutant of the *CIA2* ohnolog *CIL*, demonstrating a (partial) functional redundancy between CIA2 and CIL and that CIA2 (and therefore also CIL) shares partially overlapping functions with the two homologous barley proteins HvCMF3 and/or HvCMF7, respectively. We show that AAC proteins, representing a sub-family of the CCT domain containing proteins, act as nuclear/chloroplast proteins with a role in chloroplast development. Mutation of all four genes reduces the amount of chloroplast ribosomes, i.e., impairs chloroplast translation (mutation of *CIL* showed effects on chloroplasts only in combination with mutated *CIA2*). Impaired plastid translation leads to retrograde signaling affecting the expression of many nuclear genes, mainly those involved in the control of photosynthesis and stress response (Chan et al., 2016; Kleine and Leister, 2016; Crawford et al., 2018; Dietz et al., 2019; Zhao et al., 2020; Wu and Bock, 2021). Retrograde signaling has been analyzed in *Hvcmf7 (albostrians)* (Börner, 2017; Rotasperti et al., 2020). It needs still to be investigated if retrograde signaling may also have effects on the expression of nuclear genes in *Atcia2, Atcil* and *Hvcmf3* and affect the phenotype of the mutants. In may be interesting in this context that *cia2* and *cil* differ from the wild type in their response to certain stresses (Gawronski et al., 2020). The four AAC genes are composed of conserved domains (Figure 1) suggesting that not only AtCIL but also HvCMF3 and HvCMF7, which partially could rescue the *cia2* mutant, have function(s) similar to *CIA2,* i.e., may act as regulators of the transcription of nuclear genes involved in chloroplast biogenesis. The proposed location of HvCMF3 and HvCMF7 in the nucleus fits to this hypothesis. Interestingly, the two barley genes and possibly CIA2 are imported (also) into the chloroplasts/plastids (Li et al., 2019b; Li et al., 2019a; Gawronski et al., 2020). This observation will stimulate further research on this group of proteins, which may perform regulatory functions via protein-protein interactions not only in the nucleus (Sun et al., 2009a; Yang and Sun, 2020) but also in the plastids. The situation is reminiscent of other proteins that are dually localized in plastids and nuclei, participate in gene expression in both locations and may be involved in the communication between the organellar and nuclear genomes (Krupinska et al., 2020).

## EXPERIMENTAL PROCEDURES

### Plant Materials and Growth Conditions

Arabidopsis ecotype Col-0 was used for site-directed mutagenesis of the *CIL* gene. The plants were grown under phytochamber (poly klima, Freising, Germany) conditions with 16 h light / 8 h dark, at 20°C day / 17°C night, 65% relative humidity and photosynthetic active radiation (PAR) of 180 μmol m^-2^ s^-1^ light intensity. Screening of Cas9-induced T1 plants and progeny propagation were performed under the same phytochamber condition as described above. All the plants were grown in substrate 2 (Klasmann-Deilmann GmbH, Geeste, Germany).

Generation of the *cia2cil* double mutant: The *cia2* mutant (TAIR germplasm stock number CS6522; https://www.arabidopsis.org/servlets/TairObject?type=stock&id=1000876084) was used as maternal parent and pollinated with the *cil* mutant line AtCIL_P4_2_18_10. F1 hybrids heterozygous for the *CIA2* locus were kept for seed production. Screening for homozygous *cia2cil* double mutants was performed in the F2 generation. All plants were grown in the phytochamber as described above.

Seeds of *cil* and *cia2cil* have been deposited at NASC (Nottingham Arabidopsis Stock Center). Links to the stock page: *cil* mutant (NASC ID: N2110093), http://arabidopsis.info/StockInfo?NASC_id=2110093; *cia2cil* mutant (NASC ID: N2110094), http://arabidopsis.info/StockInfo?NASC_id=2110094.

### Vector Construction

The vectors generated in this study can be classified into three categories according to different experimental applications.

1. Vectors for site-directed mutagenesis of the *CIL* gene. To generate an RNA-guided Cas9 expression vector we first integrated a *NcoI/SpeI* fragment from pEN-Chimera (Fauser et al., 2014), containing the *U6-26* promoter from *Arabidopsis thaliana* and the gRNA-encoding chimera, into pAB-M (DNA-Cloning-Service, Hamburg, Germany), yielding in pSI55. In parallel, we generated a vector consisting of the *UBIQUITIN4-2* promoter from *Petroselinum crispum,* the *Cas9* sequence (codon optimized for *Arabidopsis thaliana*) and the *pea3A* gene from *Pisum sativum*. Therefore, the promoter and the first half of the *Cas9* sequence were amplified using pDe-Cas9 (Fauser et al., 2014) as a template and the primer pair PromCas F/PromCas R, while the second part of the Cas9 sequence together with the terminator was amplified by using the CasTerm F/CasTerm R primer combination (Supplemental Table 2). The pAB-M vector (DNA-Cloning-Service, Hamburg, Germany) was digested with *Spe*I and Gibson Assembly (NEB, Frankfurt am Main, Germany) was performed according to the manufacturer’s protocol using the beforehand amplified fragments to generate pSI56. In the last step, the gRNA-containing *Not*I/*Spe*I fragment from pSI55 was integrated into pSI56 resulting in pSI57 – the gRNA/Cas9 vector in which the gRNA-specific part (annealed oligos) can be integrated via *BbsI* restriction enzyme sites. As a result, four vectors were constructed for site-directed mutagenesis of the *CIL* gene. The derived vectors were designated as pGH502 (for PS3 target motif), pGH503 (for PS1-2 target motif), pGH505 (for PS2 target motif) and pGH508 (for PS1-1 target motif), respectively. Subsequently, the expression cassette of pGH502, pGH503, pGH505 and pGH508 was individually introduced into the binary vector p6i-d35S-TE9 (DNA-Cloning-Service, Hamburg, Germany) via the *Sfi*I restriction sites. The resulting plasmids are designated as pGH474 (for PS3 target motif), pGH475 (for PS1-2 target motif), pGH477 (for PS2 target motif) and pGH480 (for PS1-1 target motif) and were used for *Agrobacterium*-mediated transformation of ecotype Col-0.
2. Vector for Arabidopsis protoplast transformation for the determination of subcellular localization. The coding sequence of *CIL* was fused with the N terminus of the sequence encoding the GFP reporter (Chiu et al., 1996) through ligation into the *SpeI/XmaI* cloning sites of the vector pSB179 (Li et al., 2019b). The derived vector was designated as pML53 (CIL:GFP) and used for Arabidopsis protoplast transformation.
3. Vectors for genetic complementation of the *cia2* mutant. The coding sequence of *HvCMF7* and *HvCMF3* was inserted into the *SpeI/HindIII* cloning sites of pUbiAT-OCS (DNA-Cloning-Service, Hamburg, Germany) to produce pML29 and pML31, respectively. *Spe*I/*Xma*I cloning sites of pUbiAT-OCS were adopted for cloning of the truncated *CIA2* gene lacking the coding sequence for the predicted N-terminal chloroplast transit peptide (S^2^-R^59^); the derived vector is designated as pML36. Subsequently, the expression cassettes of pML29, pML31 and pML36 were individually introduced via the *Sfi*I restriction sites into the binary vector p6i-d35S-TE9 (DNA-Cloning-Service, Hamburg, Germany). The resulting plasmids are designated as pML30 (p6id35S:pUbiAT:HvCMF7), pML32 (p6id35S:pUbiAT:HvCMF3) and pML37 (p6id35S:pUbiAT:▲cTP-CIA2) and were used for *Agrobacterium-mediated* transformation of the *cia2* mutant.

### Arabidopsis Protoplast Transformation

Isolation and transformation of Arabidopsis protoplasts were performed following the protocol as described previously (Yoo et al., 2007). Briefly, protoplasts were isolated from 4-week-old Arabidopsis plants grown under the phytochamber conditions mentioned above. The protoplasts were suspended at a concentration of 2 x 10^5^ mL^-1^ in W5 solution (154 mM NaCl, 125 mM CaCl_2_, 5 mM KCl, 4 mM MES, 5 mM glucose) after counting cells under the microscope using a hemocytometer. The W5 solution was replaced by an equal volume of MMG solution (0.8 M mannitol, 1 M MgCl_2_, 100 mM MES) after incubation for 30 min on ice. Next, 100 μL of the prepared protoplasts were mixed gently with 20 μL of plasmid DNA (5 μg/μL) followed by adding and mixing completely with 110 μL of polyethylene glycol (PEG) solution (40% PEG 4000, 0.2 M mannitol, 0.1 M CaCl_2_). The transfection mixture was diluted with 400 μL of W5 solution by gently inverting the tube after keeping at room temperature for 10 min. The transfected protoplasts were collected by centrifugation at 100 *g* for 2 min., resuspended in 1 mL of WI solution (0.5 M mannitol, 20 mM KCl, 4 mM MES) and incubated in darkness at room temperature for 24 hours. Then, GFP fluorescence was checked under the laser scanning confocal microscope system LSM780 (Carl Zeiss, Jena, Germany).

### *In vitro* Chloroplast Import

The coding sequences of *CIL* and *CIA2* were cloned into pSP65 using *Bam*HI and *SalI* as restriction sites. *In vitro* transcription and translation to radiolabel the proteins with [^35^S]-methionine was performed with the TnT^®^ Quick Coupled Transcription/Translation System (Promega, Walldorf, Germany) in reticulocyte lysate. Pea plants *(Pisum sativum,* cv. ‘Arvica’) were grown on vermiculite for 10 days in a climate chamber [(14 h/10 h day-night cycle, 120 μE m^-2^s^-1^, temperatures of 20 °C/14 °C (light/dark)]. The leaf material was mixed in isolation buffer (330 mM sorbitol, 20 mM MOPS, 13 mM Tris pH 7.6, 3 mM MgCl_2_, 0.1 % BSA) filtered and centrifuged for 1 min at 1900 *g,* 4°C. The pellet was loaded on a discontinuous gradient 40% Percoll solution (330 mM sorbitol, 50 mM HEPES pH 7.6, 40% Percoll) and 80% Percoll solution (330 mM sorbitol, 50 mM HEPES pH 7.6, 80% Percoll) for 5 min at 8000 *g,* 4°C. Intact chloroplasts were washed twice with washing buffer (330 mM sorbitol, 25 mM HEPES pH 7.6, 3 mM MgCl_2_). The final pellet was resuspended in wash buffer and chlorophyll concentration was determined according to Arnon (1949). For the import reaction, 10 μg chlorophyll was used in a final reaction volume of 100 μl import buffer (300 mM sorbitol, 50 mM HEPES pH 8.0, 3 mM MgSO_2_, 50 mM ascorbic acid, 20 mM gluconate, 10 mM NaHCO3, 0.2% BSA, 4 mM MgCl_2_, 10 mM methionine, 10 mM cysteine, 3 mM ATP) together with 7 μl ^35^S-labeled, translated preprotein. Import was performed for 20 min at 25°C. 100 μl wash buffer was added and samples were centrifuged at 1500xg for 1 min, 4°C. Pellets were resuspended in SDS loading buffer and proteins were separated by a 12% SDS-PAGE, which was vacuum dried and exposed on a Phosphorimager screen for 14 h. Screens were analyzed by a Typhoon Phosphorimager (GE Healthcare, Uppsala, Sweden).

### Stable Transformation of Arabidopsis

Plasmids pML30, pML32 and pML37 were used for functional complementation of the *cia2* mutant. Vectors pGH474, pGH475, pGH477 and pGH480 were used for *Agrobacterium*-mediated transformation or co-transformation of Arabidopsis ecotype Col-0. The vectors were separately introduced into *A. tumefaciens* strain pGV2260 using a heat shock protocol (Höfgen and Willmitzer, 1988). Briefly, thaw *Agrobacterium* competent cell on ice, add 1 μg plasmids and mix gently. The mixture was incubated successively for 5 min on ice, 5 min in liquid nitrogen and 5 min at 37°C. After dilution in 1 mL LB-medium [1% (w/v) tryptone, 0.5% (w/v) yeast extract, and 1% (w/v) NaCl] the cells were shaken at 250 *rpm* for 2 hours at 28°C. Aliquots of 200 μL were plated on LB-plates [LB-medium supplemented with 0.8% (w/v) agar] containing 100 μg/mL of spectinomycin and incubated for 2 days at 28°C. Transformation of Arabidopsis was achieved by using the floral dip method (Clough and Bent, 1998). One single colony was picked from the LB-plate and incubated in 5 mL LB-medium (starter medium) with shaking at 250 *rpm* for 24 hours. Subsequently, 1 mL starter medium was diluted in 200 mL freshly prepared LB-medium and kept with shaking at 250 *rpm* for another 24 hours. The cells were resuspended in 100 mL culture medium [5% (w/v) sucrose and 0.05% (v/v) Silwet L-77] after collection by centrifugation (5500 *g*) for 10 min at room temperature. Now, the prepared infiltration medium is ready for transformation. For floral dip, infiltration medium was added to a beaker, Arabidopsis plants were inverted into the suspension such that all inflorescence tissues were submerged, and plants were then removed after 5 seconds of gentile agitation. The plants were kept in dark with 100% humidity for 24 hours and then maintained under greenhouse conditions as described above.

### Selection of Putative Transformants using an Antibiotic Marker

Arabidopsis seeds were surface sterilized with 1 mL 70% (v/v) ethanol containing 0.05% (v/v) Tween 20 (Merck, Darmstadt, Germany) by shaking on a table incubator at 1400 rpm for 3 min. After removing the supernatant, seeds were washed twice with 1mL 100% ethanol, shaked by hand and the supernatant was immediately removed. Seeds were left under a fume hood to dry for 1 hour after removing the residual ethanol. Sterilized seeds were sown on ½ MS medium [0.22% (w/v) Murashige and Skoog Basal Medium (Sigma-Aldrich M5519, Taufkirchen, Germany); 0.8% (w/v) agar; pH = 5.7] or ½ MS selection medium supplemented with 25 μg/ml hygromycin B (Thermo Fisher Scientific, Braunschweig, Germany). Plant selection was performed following the rapid method as reported (Harrison et al., 2006). In brief, seeds were stratified for 2 days in the dark at 4°C. After stratification, seeds were moved to the phytochamber (under above mentioned conditions), illuminated for 6 h in order to stimulate germination. The medium plates were then kept in dark for 2 days, wrapped with aluminium foil. The foil was removed and plates were incubated in the phytochamber for 3 days at long day conditions. Seedlings with long hypocotyl (i.e., positive transformants carrying T-DNA) were transferred into 6 cm (diameter) pots filled with substrate 2 (Klasmann-Deilmann, Geeste, Germany).

### Polymerase Chain Reaction

For mutation detection of *Cas9*-induced mutants, polymerase chain reactions (PCR) were performed in a total volume of 20 μL containing 40 ng of genomic DNA, 4 mM dNTPs, 1 μL each of 5 μM forward and reverse primers, 2 μL of 10x PCR buffer (100mM Tris-HCl, pH8.3, 500 mM KCl, 15 mM MgCl_2_, and 0.01% gelatin), and 0.5 units of HotStarTaq DNA polymerase (Qiagen, Hilden, Germany). The following touch-down PCR program was used with a GeneAmp 9700 thermal cycler (Life Technologies, Darmstadt, Germany): initial denaturation at 95°C for 5 min followed by five cycles at 94°C for 30 s, annealing at 65 to 60°C (−1°C/cycle) for 30 s, extension 1 min at 72°C, and then proceed for 40 cycles at 94°C for 30 s, 60°C for 30 s, 72°C for 1 min, and a final extension at 72°C for 10 min. For vector construction, the PCR reaction profile and program were setup as above with the following modifications: Q5 high-fidelity DNA polymerase (NEB, Frankfurt am Main, Germany) and 5x Q5 reaction buffer were used for PCR amplification. cDNA of *CIA2/CIL* (Arabidopsis ecotype Col-0)*, HvCMF3* and *HvCMF7* (barley cultivar ‘Haisa’), respectively, was used as template. cDNA synthesis was performed as described previously (Li et al., 2019b). All PCR amplicons and derived vectors were sequenced on an ABI 3730 XL DNA analyzer (Life Technologies, Darmstadt, Germany).

### Chloroplast Ultrastructural Analysis

The first leaves of the primary bolt were collected from plants used for the automated, imaging-based phenotyping experiment at developmental stage 26 DAS or DAT. For ultrastructural analysis, three biological replicates were prepared for each plant family and used for combined conventional and microwave-assisted chemical fixation, substitution and resin embedding following the protocol as described previously (Li et al., 2019a). Sectioning and transmission electron microscopy analysis was performed as described (Daghma et al., 2011).

### Determination of Chlorophyll Content

Leaf material was harvested from 25-day-old seedlings, weighted and immediately frozen in liquid nitrogen. After homogenization (Mixer Mill MM400, Retsch GmbH, Haan, Germany), 1.5 mL of N,N-dimethylformamide (DMF) was added to each sample, followed by mixing on an overhead shaker (Keison Products, Chelmsford, England) for 30 min. The supernatant obtained after centrifugation (14,000x *g* for 10 min, room temperature) was transferred to a new 2 mL Eppendorf tube. The chlorophyll content was determined according to Porra et al. (1989). In brief, cuvette-based measurement (cuvette with 1 mm path length) was conducted by help of a Spectramax Plus spectrophotometer (GENEO BioTechProducts GmbH, Germany). Chlorophyll *a* and *b* content was calculated by the following equation: chlorophyll *a* = 13.43(A^663.8^ - A^750^) - 3.47(A^646.8^ - A^750^); chlorophyll *b* = 22.90(A^646.8^ - A^750^) - 5.38(A^663.8^ - A^750^).

### High-throughput Automated, Imaging-based Phenotyping

Parameters related to photosynthetic performance were determined using an automated high throughput imaging system (Tschiersch et al., 2017). Two independent experiments were performed. Experiment I included four plant families: Col-0, *cil* mutant, *cia2* mutant and *cia2cil* mutant, with 15 plants per family. Instead of hygromycin selection, seeds of each family were directly sowed in the pots. Experiment II included five plant families: Col-0, *cia2* mutant and three families genetically complemented by *HvCMF3, HvCMF7* and ▲ *cTP_CIA2,* respectively. After antibiotic screening with hygromycin, 24 plants were selected from each of the Col-0, *cia2* mutant and *HvCMF3* families, and 96 plants were selected from each of the *HvCMF7* and ▲ *cTP_CIA2* families. The selected plants were transferred into the imaging system and phenotyping was performed following the protocols described previously (Li et al., 2019a) with the following modifications. For experiment I, measurement of PSII operating efficiency (ΦPSII) and electron transport rate (ETR) were performed for light adapted plants. Equal light intensity of 120 μE and 400 μE was independently applied during the adaptation procedure. Determination of the quenching parameters (F_v_/F_m_, NPQ, qP) were measured for dark-adapted plants with a light intensity of 120 μE. Measurements were performed at two developmental stages, 20 and 25 DAS. For experiment II, a light intensity of 180 μE was applied for determination of the chlorophyll fluorescence kinetics. Measurements were performed at 16, 18, 23 and 25 DAT.

### RNA Gel-blot Analysis

RNA isolation was performed using the TRIzol reagent (Invitrogen) following manufacturer’s instructions. One microgram of total RNA per lane was separated in 1.2% agarose/formaldehyde gels. RNA was transferred to Hybond-N (GE Healthcare, Münster, Germany) by passive transfer overnight in 25mM sodium phosphate buffer, UV cross-linked and hybridized in an Ambion^®^ ULTRAhyb^®^ at 68°C overnight with fluorescently labelled RNA probes generated by *in vitro* transcription of templates obtained by PCR using oligonucleotides as described in Supplemental Table 2. The *in vitro* transcription reaction contained 0.25 mM 5-Azido-C3-UTP (Jena Bioscience, Jena, Germany). Purified RNA probes were Click-labelled with either Cy5.5-alkyne or Cy7.5-alkyne (Lumiprobe, Hannover, Germany). Hybridized membranes were washed twice in 0.5x SSC, 0.1% (w/v) SDS and twice in 0.1x SSC, 0.1% SDS at 68°C. Membranes were scanned using the Odyssey CLx Imaging system (LI-COR, Lincoln, USA).

## SUPPLEMENTAL DATA

**Supplemental Figure 1.** Mutation detection in T1 plants by colony-PCR.

**Supplemental Figure 2.** Chloroplast import assay.

**Supplemental Figure 3.** Measurement of photosynthetic performance of wild type Col-0 and mutants *cil*, *cia2* and *cia2cil*.

**Supplemental Figure 4.** Analysis of chloroplast rRNA processing in Col-0, *cil, cia2* and *cia2cil* mutant plants.

**Supplemental Figure 5.** Quantification of ratio of mature 16S rRNA to pre-mature 16S rRNA.

**Supplemental Figure 6.** Measurement of photosynthetic performance of *cia2* mutant, complementation lines and Col-0.

**Supplemental Figure 7.** Expression profile of *CIA2* and *CIL* on eFP viewer.

**Supplemental Table 1.** Flowering time of Col-0, *cil, cia2* and *cia2cil*.

**Supplemental Table 2.** Primers used in this study.

## ACKNOWLEDGEMENTS

The authors gratefully acknowledge technical support from Mary Ziems for maintaining the Arabidopsis seeds; Susanne Koenig for Sanger sequencing; Gunda Wehrstedt and Ingo Muecke for their support in the LemnaTec experiment; Rhonda Meyer for support in cultivation of Arabidopsis; Marion Benecke and Kirsten Hoffie for microscopy; Hans-Peter Mock and Elena Brueckner for providing spectrophotometer facility and technical support on chlorophyll measurement; Cornelia Stock and Solmaz Khosravi for help in vector construction and Arabidopsis transformation; Tamara Bergius for technical assistance in the import experiments; Jochen Kumlehn for helpful discussion and supporting transformation experiments; S.S. thanks Jürgen Soll for helpful discussion. This work was supported by the Deutsche Forschungsgemeinschaft (DFG) grants STE 1102/13-1 to N.S. and CRC TR-175, project B06 to S.S.

## AUTHOR CONTRIBUTIONS

M.L., T.B. and N.S. conceived the study. M.L., H.R., M.M., A.J., H.T., G.H., S.S. and S.C. performed experiments; M.L. coordinated the project; all authors analyzed the data; M.L., T.B., and N.S. wrote the article.

